# An adaptable, user-friendly score sheet to monitor welfare in experimental fish

**DOI:** 10.1101/2023.07.19.549642

**Authors:** Mathilde Flueck-Giraud, Heike Schmidt-Posthaus, Alessandra Bergadano, Irene Adrian-Kalchhauser

## Abstract

Fish are increasingly used as experimental animals across research fields. Currently, a quarter of all experimental animals used in Europe are fish. Less than 20% of these are standard model species. Welfare assessments for experimental fish are in their infancy compared to rodents. This can be attributed to the diversity of species used, the relative recency of fish as go-to model for research, and challenges to assess welfare and pain in non-vocal underwater species. The lack of guidelines and tools presents a challenge for researchers (particularly, for newcomers), for ethics committees, and for implementing refinement measures.

Here, we present an adaptable, user-friendly score sheet for fish. The parameters contained in the excel tool are based on a literature review, have been validated by expert interviews, and evaluated by a fish pathologist. The tool allows to score individuals as well as groups, calculates summary scores and visualizes trends. We provide the underlying literature, give use examples and provide instructions on the adaptation and use of the score sheet.

We hope that this tool will empower researchers to include welfare assessment in their routines, foster discussions on fish welfare parameters among scientists, facilitate interactions with ethics committees, and most importantly, enable the refinement of fish experiments.

## Introduction

A wide range of research areas use fish as experimental models. Topics include toxicological tests ^1^, pharmacology ^2^, developmental biology ^3, 4^, complex human diseases ^5^ and behaviour ^6^. Fish are also often used to probe the response of aquatic organisms to the impact of global change, ocean warming and ocean acidification ^7^. Teleost fish share key genetic, anatomical and physiological properties with humans ^5, 8^ and therefore are valuable organisms to investigate basic evolutionary concepts ^9–12^, stem cells and regeneration ^1^^3^, the impact of chemicals and pollutants on organismic health ^14^(12), or pharmacology ^15^.

In 2020, a quarter (27.4%) of all animals used for research in the EU were fish. Zebrafish (*Danio rerio*) as the most widely known experimental fish model constituted 13% of all experimental fish ^15^, but numerous other fish species beyond zebrafish are used in research ^5, 16^, such as medaka (*Oryzias latipes*) ^17^, fat head minnow (*Pimephales promelas*) ^18^, trout (*Salmonidae*) ^19^, cichlids (*Cichlidae*) ^20^, sticklebacks (*Gasterosteidae*) ^21^, guppies (*Poecilia reticulata*) ^22^, mollies (*Poecilia latipinna*) ^23^, or gobies (*Gobidae*) ^24^, to just name a few. Evolutionarily, some of these species are more distantly related than humans to armadillos ^25^.

In accordance with widespread legal requirements to implement the 3Rs (Replace, Reduce, Refine ^26^) in animal experiments, the awareness and demand to objectively assess welfare in animal experiments continues to increase, also in research using fish. The 3Rs postulate to replace, if possible, animal models with non-living models, to reduce the number of animals necessary to fulfil an experiment, and to refine the experiments to avoid or reduce negative impacts, and they serve as foundation for laws and recommendations on animal welfare in research^27^. Measures supporting the third R, refinement, are very much dependent on the monitoring and documentation of animal welfare during husbandry and experimentation, which is legally mandatory in many countries. The tool commonly used to measure welfare are score sheets.

The aim of score sheets is to document the state of welfare and to detect alterations in welfare between groups, individuals, or time-points in a semi-quantitative, semi-objective manner. However, one can only measure what one can define and quantify, and animal welfare remains a tricky concept particularly for fish ^28^. EU regulations, for example, contain no notion of welfare ^29^. National definitions of welfare vary (ranging from the five freedoms ^30^ to dignity- or well-being-focused approaches ^31^). While fish are treated as sentient by law ^29^, and despite key initiatives ^28, 32–37^), some sources still express a need for debate on the matter ^27, 32, 38^. Knowledge on fisheś emotions and behaviour changes is available on some species ^8, 28, 39, 40^, but their subjective pain perception is sometimes discussed controversially ^32, 41^. Consequently, while the literature does discuss potential signs of distress in fish, evaluation tools for validated clinical and behavioural indicators of wellbeing are hard to come by for fish other than zebrafish ^42–44^.

In addition to these conceptual challenges, fish score sheets need to account for the previously mentioned diversity of species with distinct behaviours, morphology, physiology, and environmental requirements ^8^, and the absence of characteristics well established for rodents, such as vocalisations, body language, and spontaneous as well as induced behaviours such as grooming ^35, 45, 46^, or for agricultural species ^46^. While decipherable behaviours have been identified in some fishes, many remain understudied or uninterpretable ^40^. Fish are often housed in considerably large groups, which on the one hand complicates individual welfare monitoring and can result in specific welfare problems (e.g., hierarchy or territory issues (28)), but on the other hand expands the set of observable and measurable behavioural indicators beyond individual characteristics. If sourced from the wild, they come with a range of common background conditions ^47^.

Finally, in addition to these system-inherent challenges, training opportunities taking fish welfare into account are presently severely limited. Good welfare practices depend on skilled staff ^8, 40^, but courses for fish researchers, authorities and ethics committees are usually part of a very broad education on “non-rodent species”. Fish research publications typically do not include score sheets ^8, 28^, which limits open conversation about useful and implementable quantitative or qualitative parameters for welfare evaluations. A potential ally in experimental fish welfare is the field of aquaculture, where fish welfare translates to disease incidence, growth rates and meat quality and thus to business revenue (e.g., MyFishCheck ^48–51^, but limited mutual interactions impede the transfer of welfare related tools. Together, these challenges result in a lack of resources, expertise and frameworks with regard to fish welfare scoring in experimental research, which poses a serious challenge for researchers in the implementation of the third R for fish, particularly when researchers are new to the organism.

This publication aims to provide fish researchers with a resource facilitating the task of refinement. We provide a versatile, multispecies, modular, literature-based, and user-friendly score sheet and thereby empower researchers to assess the welfare of their animals, implement refinement strategies, and fulfil legal welfare monitoring requirements in an evidence-based manner. To develop this tool, we first systematically listed and evaluated fish welfare parameters. This entailed a scoping literature review and an exhaustive compilation of clinical and behavioural parameters that could be used as indicators of the welfare state of fish. We also conducted qualitative guided conversations with researchers and veterinarians working on different fish species and experimental systems, and reduced the list of potential indicators to a handful of individual and group parameters. Possibility of systematisation, non-intrusiveness, and modularity of parameters (e.g. module “behaviour”) were choice criteria. For each parameter, we then defined possible, easy-to-identify deviations from the desirable state, and defined mild versus severe deviations from the ideal state. We provide these parameters in the context of a modular MS office-based scoring table that a) accommodates individuals as well as groups of fish and b) can be adapted easily for the species, the experimental context or the housing situation.

We are convinced that this score sheet will simplify welfare monitoring for fish researchers worldwide, promote the use of pre-studies to optimize animal numbers and experimental conditions, facilitate 3R implementation as well as licence application procedures, and serve as useful starting point for discussions within and between labs regarding fish welfare.

## Methods

### Scoping review

To gather the published state of the art regarding fish welfare assessments, we conducted a scoping review ^52^ on welfare indicators for juvenile and adult fish. The main research question was “Which welfare indicators and score sheets are currently used for fish?”. PubMed and Web of Science were queried with 40 combinations of the terms “fish”, “rating”, “welfare”, “indicator”, “score board”, “score card”, “monitoring”, “assessment”, “humane endpoint”, “perch”, “trout”, and “zebrafish” (Supplemental File 1). When a search combination received too many hits to screen and select, the search term was refined. The relevance of papers was then assessed by screening title and the abstract. Papers were retained if these suggested the paper could contain information about fish welfare. Retained papers were then screened for welfare indicators and retained if they mentioned such parameters. Reference sections of retained publications were additionally screened for publications that did not come up in the original search and were subjected to the same screening process. 84 documents were directly retrieved from the databases in round 1. An additional 15 relevant documents were identified from snowballing the references of the round 1 documents. From the 99 documents containing welfare parameters that were retained (Supplemental File 2), a list of welfare parameters was compiled. In total, the literature search yielded 68 potential parameters (Supplemental File 2).

### Expert interviews

To understand practical considerations and the diversity of needs and implementation situations, ten fish facilities in Switzerland housing zebrafish, perch, trout, stickleback, and cichlids for research on RNA, epigenetics, genetics and genomics, development, health and infection biology, and behaviour were visited. The persons in charge of running the facility were qualitatively interviewed in semi-structured interviews with an interview guide. Questions included fish species, husbandry parameters, types of experiments done, current approaches to measure welfare (tool, criteria, frequency, group or individual level, time, responsible person, treatment of transgenic lines, reasons to not use a score sheet if none was used, source of the tool), wishes and thoughts towards score sheets (what would make it easy to use, digital or analog), and considerations concerning the collected parameters (relevance of parameters, additional suggestions, species-specifics, how to group parameters, potential endpoints, relevance of a total integrating score). Answers were recorded with pen and paper and later transcribed into an excel sheet. Also, 4 additional conversations were conducted with experts on animal welfare to understand what they would expect from or look for in a scoresheet. Questions are included in supplemental file 2. The responses, except for a summary and individual quotes, are not shared to maintain participant confidentiality.

### Parameters

The 68 parameters identified in the literature were narrowed down to a meaningful, feasible set of 29 parameters in a first round based by co-author HS, a diagnostic and research veterinarian and fish pathologist with 20+ years of experience in fish health and fish welfare, and further refined to 21 parameters for individuals and 10 parameters for groups based on the expert interviews and on discussions within the team (who feature experience working with zebrafish, trout, perch, and gobies, and rodents). Specifically, the following steps were taken: a) non-invasively observable parameters were selected (for example, blood cortisol levels as stress indicators require invasive measurements), b) related parameters were grouped (for example, swimming equilibrium ^53^ and buoyancy ^54^ were summarised to navigation problems), c) a verbose explanation for possible states was developed (for example, “normal for the species”, “mild weight loss/gain”, and “severe weight loss/gain” for the parameter “nutrition”), d) considerations that could affect scoring were collected (e.g., recent feeding may cause non-problematic abdominal swelling), and e) observable states were linked to numeric scores with 0 = ideal state, 1 = not ideal, 2 = observe, act if possible, and 3 = act immediately, potentially terminate (Table 1, see Supplemental File 3 for additional literature and species information).

**Table 1.** Parameters. Welfare parameters, explanations of these parameters, and signs to look for. Includes scores in numbers and in verbose expressions. This table is translated into an interactive scoring tool in Supplemental File 4.

## Results

### Expert Interviews

Expert interviews confirmed the idea that fish welfare assessments and their use in refinement are an area of development with unsatisfactory options for fish researchers. A frequent practice was to copy other researcher’s score sheets and to satisfy administrative requests (“Once the canton of … requested more parameters, so we just added one of them”). Opinions on welfare were often based on years of experience working with fish, rather than on explicitly scorable, model specific, and objective criteria. Only 3 of the 10 facilities indicate using a score sheet as welfare tool. Two facilities stated not using a score sheet because it was not compulsory in their region. Usually, more than 1 person was in charge of assessing fish health and state at the same time. Some facilities stated that certain types of physical/morphological aberrations or behavioral signs were considered as induced by standard lab conditions and were therefore currently not considered in their score sheets as welfare issues.

With regard to a score sheet, isolated parameters were generally deemed insufficient to conclude on the welfare state of individuals and groups. Also, behavioral signs of impairment were considered more relevant than physical signs, which may be linked to the current lack of understanding regarding the relevance of e.g., injury and related pain for fish wellbeing. The relevance of an integrative total score was generally supported as reasonable given the mutual impacts of parameters (e.g., damaged fins might lead to altered swimming behavior). A request was to include potential measures to alleviate the identified issues in a score sheet, since euthanasia of impacted fish would often create issues for experimental design. A digital version was preferred, and a tablet-compatible app was suggested by several participants.

### Score sheet

The score sheet developed based on the review and the interviews is available as Supplemental File 4. It contains 21 potentially relevant parameters for individuals, and 10 parameters for groups. These 21 parameters are based on literature from 17 distinct fish species from 11 taxonomic families. They are classified into the categories body condition, skin, fins, eyes, gills/opercula, infection, swimming behavior, feeding behavior for individuals, and 2 additional categories (group behavior and mortality) for groups.

For individually housed or marked fish, for example in cichlid behaviour experiments ^6^, the status for a certain parameter is assessed verbosely by choosing from a drop-down menu (Figure 1). For example, for fin integrity, researchers can choose between “fin in healthy condition”, “slightly eroded / slightly missing”, “moderate fin loss” or “severe fin loss / missing fin”. The sheet then automatically calculates a numerical score. For fish housed in groups, which is for example usually the case for zebrafish, the estimated percentage of fish in the tank with certain issues is recorded (Figure 1). For example, 80% of fish may be scored, for a particular parameter, to display “no abnormalities / lesions”, 10% with “slight abnormalities / lesions”, 5% with “moderate abnormalities / lesions” and 5% with “severe abnormalities / lesions”. A weighted average is then calculated for the tank based on formulas already implemented in the score sheet. If percentages do not sum up to 100 due to entry errors, the respective cells in the excel sheet turn orange.

**Figure 1.**
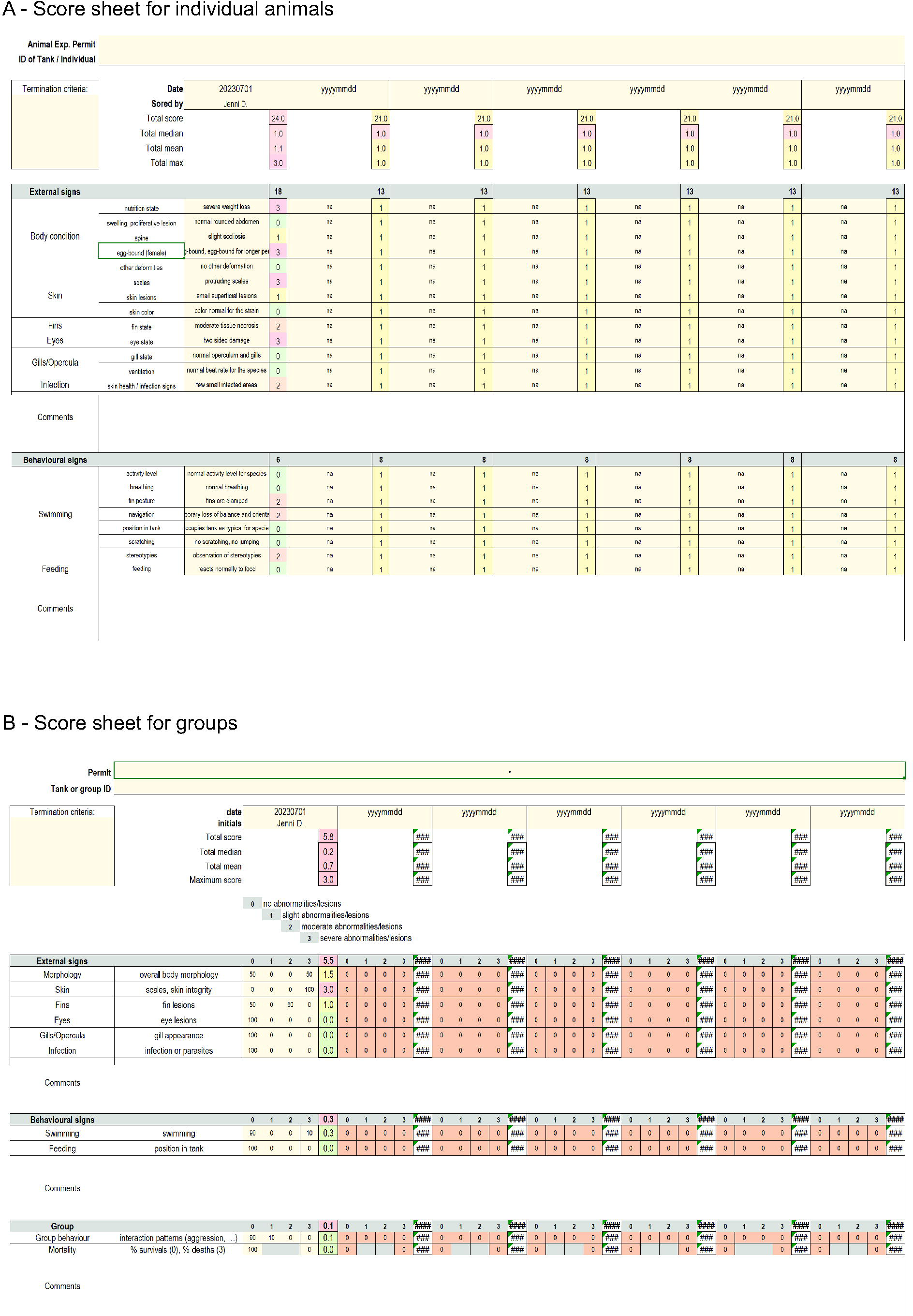
Score sheets. **A –** Score sheet for individual animals. **B** – Score sheet for groups of animals. These are screenshots taken from the interactive scoring tool provided as Supplemental File 4.

In the score sheet for individually housed or marked fish, skipping the assessment of a parameter in the individual score sheet results in a default score of 1 to avoid the false assumption of good welfare based on non-scoring. If parameters cannot be scored because it is either not feasible or not relevant regarding the circumstances, we recommend adapting the score sheet and deleting the parameter (s) (Supplemental File 4 features instructions on how to do that).

The score sheet tool provides a total score, a median, mean and maximum score for each assessment. Both these integrative and the individual scores are color coded from green to red on a relative scale for easy visual overview. 7 assessments (e.g., for one week) are designed to fit on an A4 printout for easy collection and recording of past assessments. In addition, a trend tab allows the tracking and visualization of integrative and individual scores across time. To visualize trends, users need to copy-paste values (without formulas) from areas outlined with heavy borders from the score sheet tab to the trends tab. Up to 31 data points (e.g., for one month) can be accommodated within the current dimensions of the trend tab.

#### Use scenarios

Below, we use three hypothetical common research scenarios, to demonstrate how the score sheet can be used to monitor welfare or refine experimental conditions. These scenarios are 1) a pre-trial to identify useful welfare scoring parameters for a genetically modified zebrafish strain with morphological impairments, 2) a pretrial to define a suitable acclimatization period of wild-caught individuals before an experiment, and 3) the monitoring of fish welfare during an infection trial to identify the appropriate termination point.

In scenario 1 (Figure 2), a strain of genetically modified zebrafish with developmental morphological defects in spine and eyes, and slightly elevated mortality, is to be included in a chemical exposure experiment (similar to e.g. ^55^. The researchers are confronted with the task to identify welfare parameters that are suited, in this impaired strain, to detect any additional impairment caused by the experiment. To start with, they score a control tank over a month using the tool for fish that are housed in groups. Scores relating to anatomy (morphology and eye scores) are constitutively high due to the developmental impairment. Swimming behavior is also impacted by the strains’ defects, resulting in overall high behavioral scores. Skin parameters and feeding behavior fluctuate slightly during the observation time. Mortality scores indicate that strain mortality is 10% higher than usually observed in the wildtype. Based on these documented and quantified observations, the researchers come to the following conclusions for their main exposure experiment: they will pay particular attention to skin and feeding parameters, since these parameters display variation already without treatment, and may therefore be sensitive to stressors. Also, they take the 10% mortality rate into account during sample size and power calculations and start the experiment with 10% more animals than required to achieve the desired power. They design an experiment-specific score sheet where they record anatomy and swimming behavior but – given that these parameters are high to begin with, reducing their sensitivity – flag them as potentially unhelpful. They also use the data to justify animal numbers in their animal experimentation permit application.

**Figure 2.**
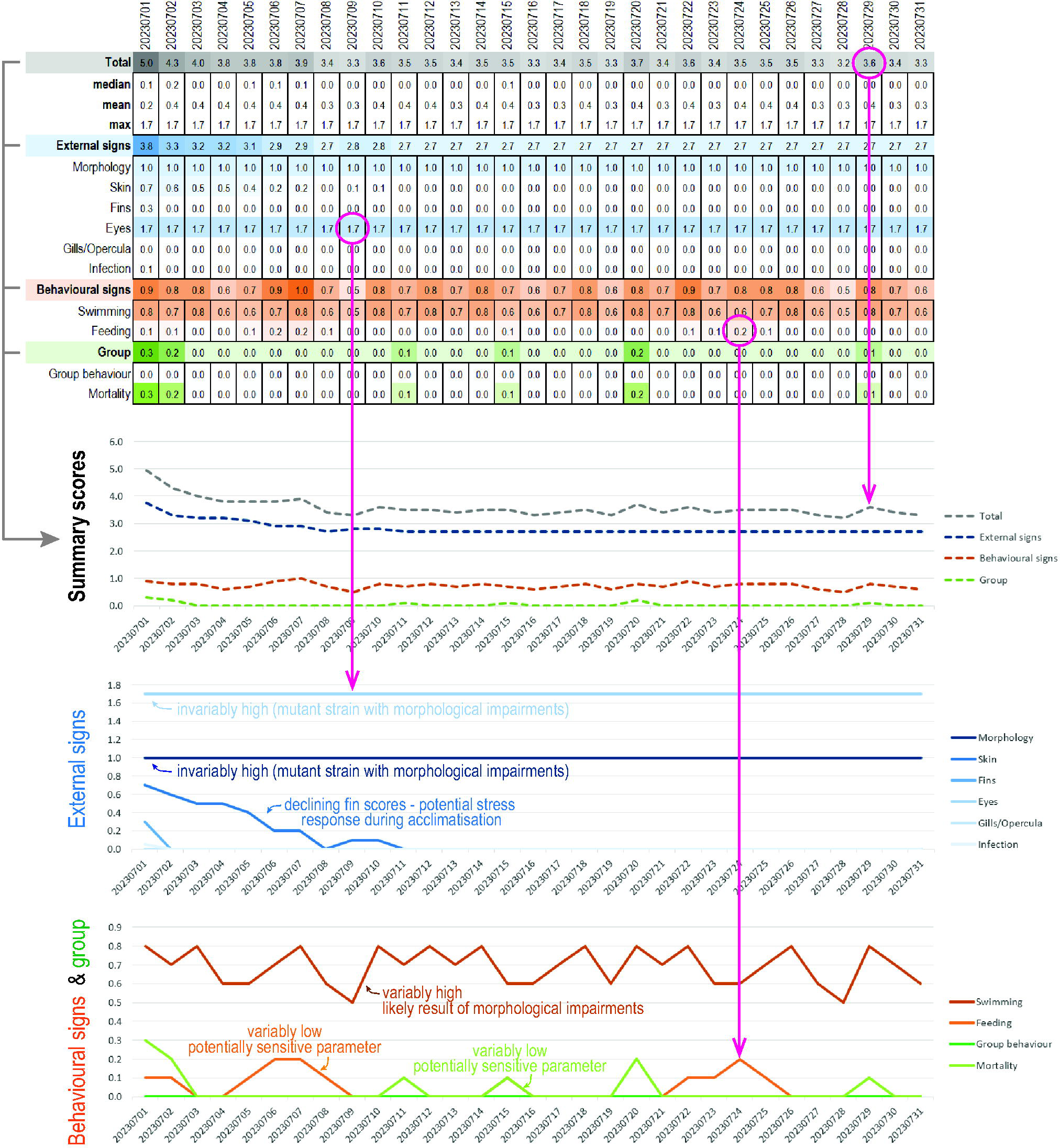
Application Scenario 1. Example for a group scoring of an impaired mutant line of zebrafish. View of the trends tab. The trends tab translates the scores into visual patterns, on the one hand through score-dependent coloring of the cells, on the other hand through trend lines. Each value in the top table corresponds to a datapoint in the trend lines (some pointed out by circles and arrows). The summary scores, external signs, and group/behavioural signs are each collated in a separate trend line figure.

In scenario 2 (Figure 3), individuals from a wild fish species are caught and brought to the lab for a behavior experiment (choice between two compartments, e.g. ^56^) before being released again. The researchers are confronted with the task to determine a suitable acclimation period before the experiment: not too long to avoid unnecessary long captivity, not too short to avoid confounding impacts of the stress during capture on the experiment. They catch several fish for a pre-trial and assess their welfare individually for a month. Figure 3 shows data from one representative individual. During the entire observation period, no signs of external damage, lesion or infection are observed. However, a diverse range behavioral abnormalities (e.g., stereotypical swimming and startle responses to movement and sounds) are observed between d1 and d13. These reactions decline in intensity and disappear in week 3. From d19 onwards, no behavioral abnormalities are observed. Based on the pretrial, the researchers therefore conclude (and have data to show) that an 18 day acclimation period is both required and sufficient for the planned behavioral experiments, and choose their setup accordingly to ensure the reliability and validity of their data.

**Figure 3.**
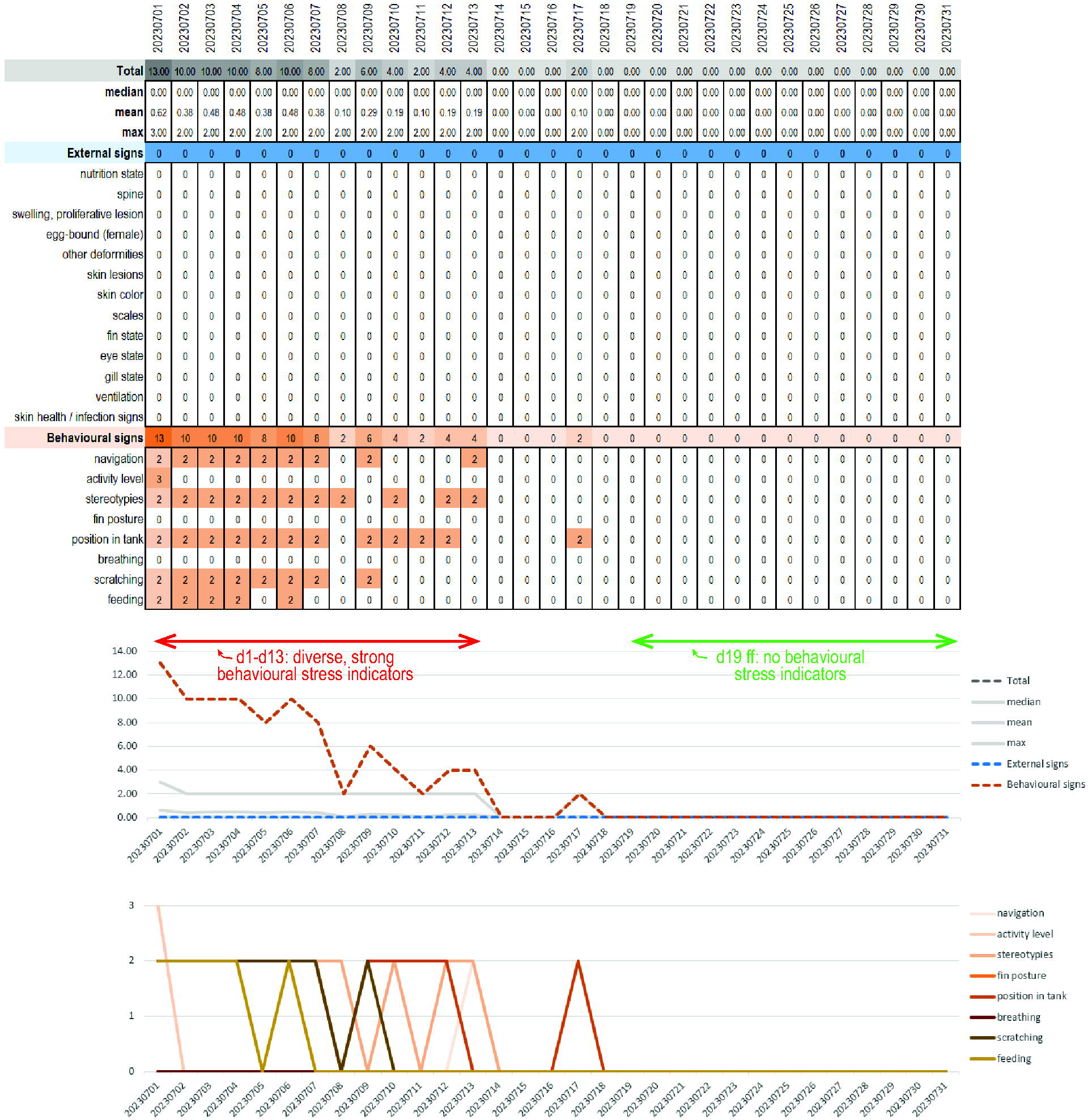
Application Scenario 2. Example for an individual scoring of a wild-caught fish during the acclimatization period. View of the trends tab. Both the score-dependent coloring on the top and the trend lines at the bottom suggest that the fish is stressed until day 13 of the acclimatization. No behavioural signs of stress are observed after day 19.

In scenario 3 (Figure 4), researchers want to perform a drug screen on zebrafish infected with a certain pathogen (similar to e.g. ^57^) to find compounds that are active against said pathogen. The researchers are confronted with the task of determining an experimental duration that is long enough to show differences between control and treatment, and short enough to avoid unnecessary suffering. They also need to determine a termination score (i.e. a cutoff where a sick animal is euthanized and removed from the experiment to avoid excessive suffering). A group of 10 fish is therefore used for a 4-week pretrial under control conditions (pathogen infection without drug) and scored with the group tool. The score sheet reveals that first skin defects are observed 14 days after pathogen exposure, and continuously increase in severity. On day 23 after pathogen exposure, swimming issues (loss of buoyancy) are starting to be observed and increase in prevalence until the end of the pretrial. Based on this pretrial, the researchers determine to run the drug screen for 25 days, since meaningful differences between control and treatment should be observable by this timepoint without causing unnecessary suffering of experimental animals. They also decide that they will terminate any tank x compound combination where the total score exceeds 11 two days in a row, since this score should not be reached in the control within 25 days and would indicate that the compound causes harm.

**Figure 4.**
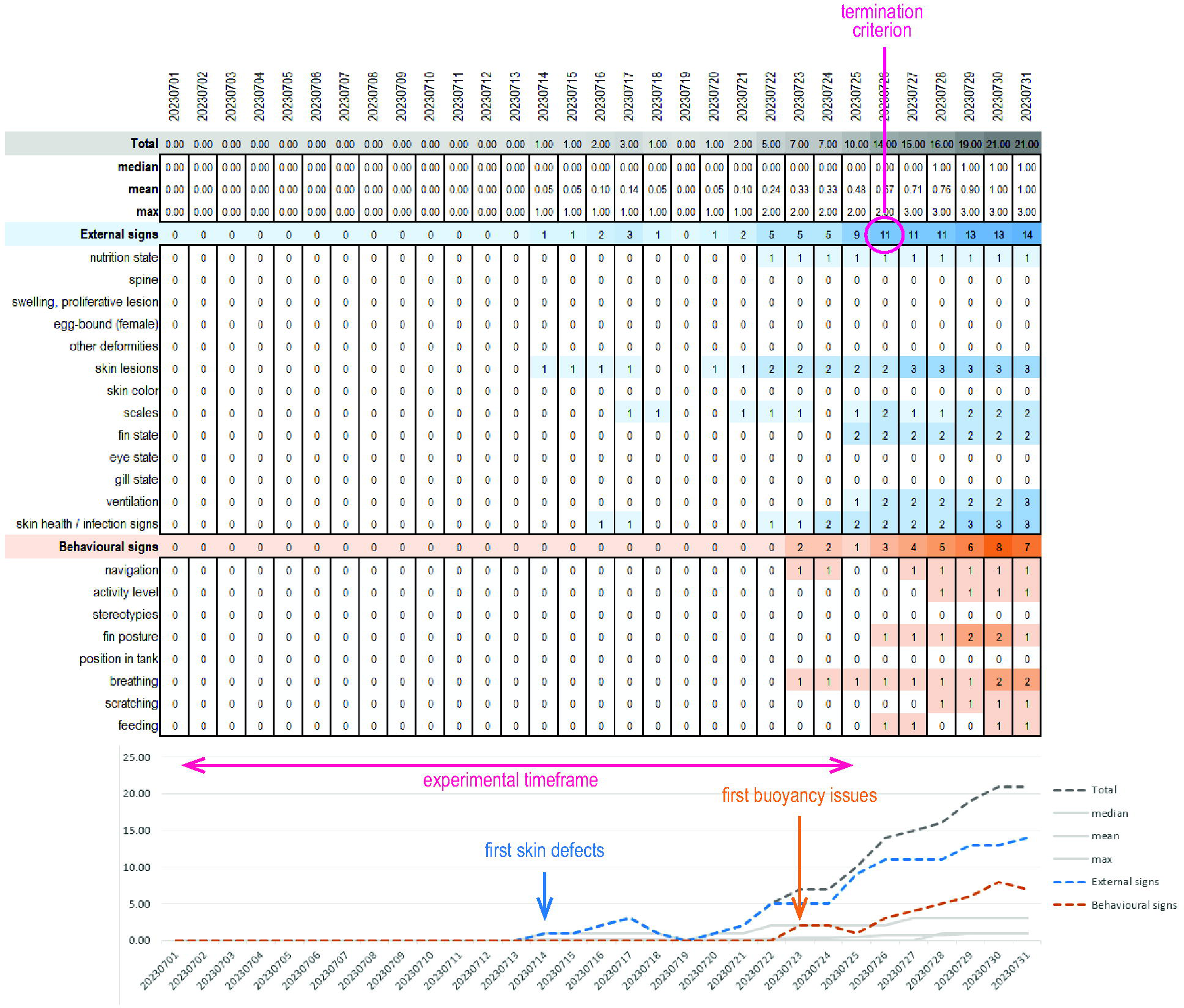
Application Scenario 3. Example for an individual scoring of an experimental fish exposed to a compound. External signs (i.e. skin defects) are observed before the appearance of behavioural signs and get progressively worse over the course of several weeks. Based on these data, the end point of the planned experiment is determined after onset of signs, but before welfare is too compromised.

#### Use and adaptation of the score sheet

Scoring 21 parameters on multiple fish every day is clearly not a feasible task. We want to emphasize that a key step in the use of this (and any other) score sheet is the adaptation phase. This phase includes a) test scoring of the parameters, followed by b) a choice of parameters that are truly useful to assess welfare in the given species, situation, and experiment, and c) creating an experiment-specific score sheet by deleting non-informative rows from the score sheet, adding parameters, or adapting the weighting.

We strongly encourage a rigorous and information-driven reduction of the score sheet to achieve maximum information and maximum compliance with minimal effort. The score sheet file contains instructions to help with the adaptation and implementation phase (Supplemental File 4, tab “instructions”). The adaptation phase encompasses 7 steps: “Adapt to setting”, “Get familiar”, “Generate baseline”, “Adapt criteria and weights”, “Termination criteria”. We hope to discourage uncritical use of the score sheet with these instructions, since this would most likely lead to frustration and non-compliance. In addition, the score sheet file contains instructions for use after adaptation. The use phase is based on three steps: “Data entry”, “Documentation”, “Statistics”. The aim is to encourage sustainable long-term preservation strategies and support good data management practices, e.g. for reporting to ethics committees.

## Discussion & Conclusion

This publication aims to provide a flexible framework to develop suited score sheets across a number of species, experimental scenarios, and housing conditions. This is by no means the first attempt to develop score sheets for experimental fish (see, for example, a zebrafish-specific pain and distress score sheet in ^58^ Table 3). However, to our knowledge, it is the first attempt to create an adaptable tool that is explicitly not supposed to be used as is, but rather to be molded by the researcher to really fit their needs regarding welfare monitoring.

### Applicability

The diversity of fish species kept for research purposes, and their housing both as single fish and large groups is a challenge for welfare assessment. The presented work tackles this in two ways. First, the developed score sheet remains adaptable. Parameters can be removed (which is actually recommended), changed, or added (e.g., for highly specific behaviours, such as mouth brooding or nest building). Second, the score sheet emphasizes the importance of relative assessments. It does not provide absolute numerical cutoffs a priori but empowers users to develop their own meaningful termination criteria based on the respective species, strain, facility, and housing conditions.

The operationalization of termination criteria (endpoints where an experiment must be terminated and the animals are euthanized to prevent further suffering) is one of the most important aspects and purposes of score sheets. How to arrive at a combined measure of maximally tolerable burden is one of the most important, but also most difficult and controversial parts of welfare assessments. Importantly, termination criteria depend heavily on the weighing of interests and the purpose of the experiment (which extent of suffering is justifiable for the expected gain of knowledge). This score sheet should facilitate the definition of termination criteria and facilitate conversations on termination criteria among fish researchers.

### Implications

This score sheet should benefitfish and fish researchers in five distinct ways. Firstly, the tool aims to empower the individual researcher. Our interviews revealed a prevalence of subjective assessments which may or may not reflect animal welfare criteria, a certain acceptance of stereotypies as given, and a dependence on few knowledgeable care takers. All these aspects could benefit from more objective scoring options. Secondly, the tool potentially fosters within-group interactions and cross-facility communication regarding welfare and termination criteria. This may improve collective welfare knowledge in the fish research community and inspire initiatives and research towards better welfare assessment in fish. For example, our interviews revealed that standard laboratory conditions are generally assumed to guarantee good welfare. One way to assess this would be to conduct a project using the tool across facilities and housing conditions. Thirdly, the tool may facilitate regulatory processes surrounding animal experimentation. In Switzerland, score sheets are required to obtain animal experimentation permits. This tool may thus support researchers in defining and submitting evidence-based monitoring and stop-criteria. Fourthly, ethics committees are usually not familiar with welfare of fish. This tool should facilitate the work of ethics committees to support well-planned and well-monitored experimentation with fish, and the communication between researchers and committees. A fifth aspect relates to teaching. Score sheets are valuable tools to discuss fish needs, health, and welfare, and can be used to train junior researchers in accurate and attentive evaluation of signs and behaviours. For example, a course could entail the application of a score sheet to strains with known physical and behavioral impairments, or to a range of species.

### Limitations

From the practical perspective, welfare monitoring can pose a challenge regarding the time requirements and personnel resources of research groups. We strongly recommend prioritizing the monitoring of few meaningful parameters at longer time intervals over more complex monitoring plans which fail in the long-term. A key factor in welfare monitoring is compliance and constancy, and these need to be given priority over perfection due to overburdening the researchers or animal care takers. From the conceptual perspective, discussions on animal welfare are often impacted by debates on fish nociception. We support the point of view that the presence of protective responses and avoidance learning may be sufficient indicators of pain, health, and welfare in fish, but agree that scoring of pain remains a challenge.

### Recommendations

In summary, this manuscript provides an objective, adaptable welfare score sheet tool to the wider fish research communityallowing monitoring of fish welfare in a semi-objective manner Prospectively, it would be valuable to engage in cross-species and cross-field efforts to build on the presented monitoring tool. Also the development of an adaptable welfare-monitoring smart-phone app would be extremely practical for the daily monitoring. Also, visual tracking applications ^59^ and sensors have great potential to facilitate fish welfare monitoring in the future.

## Acknowledgements

We are grateful to numerous experts who contributed their time and knowledge in personal conversations and during site visits, in particular (in alphabetical order) Heinz Belting, Fabienne Chabaud, Nicolas Diserens, Anna Gliva, Marcel Häsler, Ahmet Kürk, Ines Dos Santos, Alba Aparicia Fernandez, Sebastian Leidel, Nadia Mercader, Catherine Peichel, Verena Saladin, Dragan Stajic, Attila Rüegg, Jaques Voland, Gilles Willemin, and Hanno Würbel. We thank Catherine Peichel, Hanno Würbel, and Bernhard Völkl for their supportive and insightful feedback on earlier versions of this manuscript.

## Author contributions

IAK conceived the project, HS and IAK designed the analysis with input from MFG, MFG collected the data and performed the analyses with support from AB and HS, HS and IAK supervised and guided analyses, MFG wrote the manuscript draft, HS, IAK, AB and MFG edited the manuscript.

## Research ethics

No animals were used for this research.

## References

1. DeFoe D, Drummond R, Geiger D, et al. *Behavioral and Morphological Changes in Fathead Minnow (Pimephales promelas) as Diagnostic Endpoints for Screening Chemicals According to Mode of Action*. West Conshohocken, Pa.: ASTM International, 1986.

2. Bolis CL, Piccolella M, Dalla Valle AZ, et al. Fish as model in pharmacological and biological research. Pharmacological Research 2001; 44: 265–280, https://www.sciencedirect.com/science/article/pii/S104366180190845X (2001).

3. Marques IJ, Ernst A, Arora P, et al. Wt1 transcription factor impairs cardiomyocyte specification and drives a phenotypic switch from myocardium to epicardium. Development 2022; 149.

4. Kratochwil CF, Sefton MM and Meyer A. Embryonic and larval development in the Midas cichlid fish species flock (Amphilophus spp.): a new evo-devo model for the investigation of adaptive novelties and species differences. BMC Dev Biol 2015; 15: 12.

5. Schartl M. Beyond the zebrafish: diverse fish species for modeling human disease. Dis Model Mech 2013; 7: 181–192.

6. Kasper C, Kölliker M, Postma E, et al. Consistent cooperation in a cichlid fish is caused by maternal and developmental effects rather than heritable genetic variation. Proc Biol Sci 2017; 284.

7. Clark TD, Raby GD, Roche DG, et al. Ocean acidification does not impair the behaviour of coral reef fishes. Nature 2020; 577: 370–375.

8. Toni M, Manciocco A, Angiulli E, et al. Review: Assessing fish welfare in research and aquaculture, with a focus on European directives. Animal 2019; 13: 161–170.

9. Peichel CL and Marques DA. The genetic and molecular architecture of phenotypic diversity in sticklebacks. Philos Trans R Soc Lond B Biol Sci 2017; 372.

10. Liu Z, Roesti M, Marques D, et al. Chromosomal Fusions Facilitate Adaptation to Divergent Environments in Threespine Stickleback. Mol Biol Evol 2022; 39.

11. Archambeault SL, Bärtschi LR, Merminod AD, et al. Adaptation via pleiotropy and linkage: Association mapping reveals a complex genetic architecture within the stickleback Eda locus. Evol Lett 2020; 4: 282–301.

12. Ishikawa A, Kabeya N, Ikeya K, et al. A key metabolic gene for recurrent freshwater colonization and radiation in fishes. Science 2019; 364: 886–889.

13. Marques IJ, Sanz-Morejón A and Mercader N. Ventricular Cryoinjury as a Model to Study Heart Regeneration in Zebrafish. In: Poss KD and Kühn B (eds) Cardiac Regeneration: Methods and Protocols. New York, NY: Springer US, 2021, pp. 51– 62.

14. Segner H. Zebrafish (Danio rerio) as a model organism for investigating endocrine disruption. Comparative Biochemistry and Physiology Part C: Toxicology & Pharmacology 2009; 149: 187–195, https://www.sciencedirect.com/science/article/pii/S153204560800197X (2009).

15. European commission. ALURES: Section 2 -Details of all uses of animals for research, testing, routine production and education and training purposes in the EU, https://webgate.ec.europa.eu/envdataportal/content/alures/section2_number-of-uses.html (2023).

16. Mocho J-P and Krogh K von. A FELASA Working Group Survey on Fish Species Used for Research, Methods of Euthanasia, Health Monitoring, and Biosecurity in Europe, North America, and Oceania. Biology (Basel) 2022; 11.

17. Kawasaki T, Saito K, Deguchi T, et al. Pharmacological characterization of isoproterenol-treated medaka fish. Pharmacological Research 2008; 58: 348–355, https://www.sciencedirect.com/science/article/pii/S1043661808001758 (2008).

18. Parrott JL, Alaee M, Wang D, et al. Fathead minnow (Pimephales promelas) embryo to adult exposure to decamethylcyclopentasiloxane (D5). Chemosphere 2013; 93: 813–818.

19. Schmidt-Posthaus H, Ros A, Hirschi R, et al. Comparative study of proliferative kidney disease in grayling Thymallus thymallus and brown trout Salmo trutta fario: an exposure experiment. Dis Aquat Organ 2017; 123: 193–203.

20. Jadhao AG and Malz CR. Nicotinamide adenine dinucleotide phosphate (NADPH)-diaphorase activity in the brain of a cichlid fish, with remarkable findings in the entopeduncular nucleus: a histochemical study. J Chem Neuroanat 2004; 27: 75–86.

21. Wegner KM, Kalbe M, Rauch G, et al. Genetic variation in MHC class II expression and interactions with MHC sequence polymorphism in three-spined sticklebacks. Mol Ecol 2006; 15: 1153–1164.

22. Li M-H. Effects of nonylphenol on cholinesterase and carboxylesterase activities in male guppies (Poecilia reticulata). Ecotoxicol Environ Saf 2008; 71: 781–786.

23. Ayllon F and Garcia-Vazquez E. Induction of micronuclei and other nuclear abnormalities in European minnow Phoxinus phoxinus and mollie Poecilia latipinna: an assessment of the fish micronucleus test. Mutation Research/Genetic Toxicology and Environmental Mutagenesis 2000; 467: 177–186, https://www.sciencedirect.com/science/article/pii/S1383571800000334 (2000).

24. Adrian-Kalchhauser I, Hirsch PE, Behrmann-Godel J, et al. The invasive bighead goby Ponticola kessleri displays large-scale genetic similarities and small-scale genetic differentiation in relation to shipping patterns. Mol Ecol 2016; 25: 1925– 1943.

25. Meadows JRS and Lindblad-Toh K. Dissecting evolution and disease using comparative vertebrate genomics. Nature Reviews Genetics 2017; 18: 624–636.

26. Russel WMS and Burch RL. *The Principles of humane experimental technique* by W.M. Russel and R.L. Burch. London: Methuen, 1959.

27. Aleström P, D’Angelo L, Midtlyng PJ, et al. Zebrafish: Housing and husbandry recommendations. Lab Anim 2020; 54: 213–224.

28. Johansen R, Needham JR, Colquhoun DJ, et al. Guidelines for health and welfare monitoring of fish used in research. Lab Anim 2006; 40: 323–340.

29. European Parliament. Directive 2010/63/EU: LEX-FAOC098296, 2010.

30. Brambell FWR. Report of the Technical Committee to enquire into the Welfare of Animals kept under Intensive Livestock Husbandry Systems, etc. [Chairman, Professor F.W. Rogers Brambell.] London, 1965.

31. Bundesversammlung der Schweizerischen eidgenossenschaft. Tierschutzgesetz Art.3: TSchG, 2022.

32. Lambert H, Cornish A, Elwin A, et al. A Kettle of Fish: A Review of the Scientific Literature for Evidence of Fish Sentience. Animals 2022; 12.

33. Sneddon LU. Pain perception in fish: indicators and endpoints. ILAR J 2009; 50: 338–342.

34. Sneddon LU. Pain in aquatic animals. J Exp Biol 2015; 218: 967–976.

35. Sneddon LU. Evolution of nociception and pain: evidence from fish models. Philos Trans R Soc Lond B Biol Sci 2019; 374: 20190290.

36. Sneddon LU, Braithwaite VA and Gentle MJ. Do fishes have nociceptors? Evidence for the evolution of a vertebrate sensory system. Proc Biol Sci 2003; 270: 1115– 1121.

37. Griffin G and Gauthier C. Guidelines development and scientific uncertainty: use of previous case studies to promote efficient production of guidelines on the care and use of fish in research, teaching and testing. Animal Welfare 2004; 13: S181–S186, https://www.cambridge.org/core/article/guidelines-development-and-scientific-uncertainty-use-of-previous-case-studies-to-promote-efficient-production-of-guidelines-on-the-care-and-use-of-fish-in-research-teaching-and-testing/2BEA30019C80AF0EC14C953F2440472E (2004).

38. Browman HI, Cooke SJ, Cowx IG, et al. Welfare of aquatic animals: where things are, where they are going, and what it means for research, aquaculture, recreational angling, and commercial fishing. ICES Journal of Marine Science 2019; 76: 82–92.

39. Jarvis S, Ellis MA, Turnbull JF, et al. Qualitative Behavioral Assessment in Juvenile Farmed Atlantic Salmon (Salmo salar): Potential for On-Farm Welfare Assessment. Front Vet Sci 2021; 8: 702783.

40. Reilly SC, Quinn JP, Cossins AR, et al. Behavioural analysis of a nociceptive event in fish: Comparisons between three species demonstrate specific responses. Applied Animal Behaviour Science 2008; 114: 248–259.

41. Volpato GL. Challenges in assessing fish welfare. ILAR J 2009; 50: 329–337.

42. Goodwin N, Karp NA, Blackledge S, et al. Standardized Welfare Terms for the Zebrafish Community. Zebrafish 2016; 13 Suppl 1: S164–8.

43. OECD. OECD guideline for testing of chemicals.

44. Reed B and Jennings M. Guidance on the housing and care of zebrafish, 11–2010.

45. Brown C. Fish intelligence, sentience and ethics. Anim Cogn 2015; 18: 1–17.

46. Dalmau A, Velarde A, Scott K, et al. Welfare Quality® assessment for pigs (sows and piglets, growing and finishing pigs), 2009.

47. Mangus LM, França MS, Shivaprasad HL, et al. Research-Relevant Background Lesions and Conditions in Common Avian and Aquatic Species. ILAR J 2021; 62: 169–202.

48. Tschirren L, Bachmann D, Güler AC, et al. MyFishCheck: A Model to Assess Fish Welfare in Aquaculture. Animals 2021; 11.

49. Browning H. Improving welfare assessment in aquaculture. Front Vet Sci 2023; 10.

50. Garcia de Leaniz C, Gutierrez Rabadan C, Barrento SI, et al. Addressing the welfare needs of farmed lumpfish: Knowledge gaps, challenges and solutions. Reviews in Aquaculture 2022; 14: 139–155.

51. Noble C, Gismervik K, Iversen M, et al. Welfare Indicators for farmed Atlantic salmon: Tools for assessing fish welfare, nofinma, 2018.

52. Munn Z, Peters MDJ, Stern C, et al. Systematic review or scoping review? Guidance for authors when choosing between a systematic or scoping review approach. BMC Med Res Methodol 2018; 18: 143.

53. Saxby A, Adams L, Snellgrove D, et al. The effect of group size on the behaviour and welfare of four fish species commonly kept in home aquaria. Applied Animal Behaviour Science 2010; 125: 195–205.

54. Borges AC, Pereira N, Franco M, et al. Implementation of a Zebrafish Health Program in a Research Facility: A 4-Year Retrospective Study. Zebrafish 2016; 13 Suppl 1: S115–26.

55. Baraban SC, Dinday MT and Hortopan GA. Drug screening in Scn1a zebrafish mutant identifies clemizole as a potential Dravet syndrome treatment. Nat Commun 2013; 4: 2410.

56. Jutfelt F, Sundin J, Raby GD, et al. TwolJcurrent choice flumes for testing avoidance and preference in aquatic animals. Methods Ecol Evol 2017; 8: 379–390.

57. Kam JY, Hortle E, Krogman E, et al. Rough and smooth variants of Mycobacterium abscessus are differentially controlled by host immunity during chronic infection of adult zebrafish. Nat Commun 2022; 13: 952.

58. Martins T, Valentim AM, Pereira N, et al. Anaesthesia and analgesia in laboratory adult zebrafish: a question of refinement. Lab Anim 2016; 50: 476–488.

59. Quevedo C, Behl M, Ryan K, et al. Detection and Prioritization of Developmentally Neurotoxic and/or Neurotoxic Compounds Using Zebrafish. Toxicol Sci 2019; 168: 225–240.

